# A specific sequence in the genome of respiratory syncytial virus regulates the generation of copy-back defective viral genomes

**DOI:** 10.1101/349001

**Authors:** Yan Sun, Eun Ji Kim, Sébastien A. Felt, Louis J. Taylor, Divyansh Agarwal, Gregory R. Grant, Carolina B. López

**Author notes:** **Corresponding author:** Carolina B. López, PhD. Department of Pathobiology, School of Veterinary Medicine, University of Pennsylvania, Philadelphia, PA 19104, United States.

## Abstract

Defective viral genomes of the copy-back type (cbDVGs) are the primary initiators of the antiviral immune response during infection with respiratory syncytial virus (RSV) both *in vitro* and *in vivo.* However, the mechanism governing cbDVG generation remains unknown, thereby limiting our ability to manipulate cbDVG content in order to modulate the host response to infection. Here we report a specific genomic signal that mediates the generation of RSV cbDVGs. Using a customized bioinformatics tool, we identified regions in the RSV genome frequently used to generate cbDVGs during infection. We then created a minigenome system to validate the function of one of these sequences and to determine if specific nucleotides were essential for cbDVG generation at that position. Further, we created a recombinant virus that selectively produced a specific cbDVG based on variations introduced in this sequence. The identified sequence was also found as a common site for cbDVG generation during natural RSV infections, and common cbDVGs generated at this sequence were found among samples from various infected patients. These data demonstrate that sequences encoded in the viral genome are critical determinants of the location of cbDVG generation and, therefore, this is not a stochastic process. Most importantly, these findings open the possibility of genetically manipulating cbDVG formation to modulate infection outcome.

**Author summary:** Copy-back defective viral genomes (cbDVGs) regulate infection and pathogenesis of *Mononegavirales.* cbDVG are believed to arise from random errors that occur during virus replication and the predominant hypothesis is that the viral polymerase is the main driver of cbDVG generation. Here we describe a specific genomic sequence in the RSV genome that is necessary for the generation of a large proportion of the cbDVG population present during infection. We identified specific nucleotides that when modified altered cbDVG generation at this position, and we created a recombinant virus that selectively produced cbDVGs based on mutations in this sequence. These data demonstrate that the generation of RSV cbDVGs is regulated by specific viral sequences and that these sequences can be manipulated to alter the content and quality of cbDVG generated during infection.

## Introduction

Defective viral genomes (DVGs), which are generated during the replication of most RNA viruses, potentiate the host innate immune response [1-5] and attenuate the infection *in vitro* and *in vivo* [4, 6-9]. Importantly, in naturally infected humans, the presence of DVGs correlates with enhanced antiviral immune responses during RSV infection [6] and reduced disease severity in influenza virus infection [8]. Significant effort is currently invested in harnessing DVGs as antivirals due to their strong immunostimulatory activity and ability to interfere with the replication of the standard virus. However, despite over 50 years of appreciating their critical functions in multiple aspects of viral infections, the molecular mechanisms that drive DVG generation remain largely unknown. This lack of understanding hampers our ability to effectively harness DVGs for therapeutic purposes and limits our capacity to generate tools to elucidate their mechanism of action and impact during specific viral infections.

There are two major types of DVGs: deletion and copy-back (cb) [10]. Both types are unable to complete a full replication cycle without the help of a co-infecting full-length virus [11,12] and can be packaged to become part of the viral population [13]. Deletion DVGs, common in influenza virus and positive strand RNA viruses, retain the 3’ and 5’ ends of the viral genomes but carry an internal deletion [14-16]. These types of DVGs are believed to arise from recombination events [17,18] and can have a strong interfering activity over the standard virus [19]. cbDVGs are common products of negative sense (ns)RNA virus replication, including Sendai virus, measles virus, and respiratory syncytial virus (RSV), and are the primary stimulators of the innate immune response during nsRNA virus infection [6, 7, 20, 21], cbDVGs arise when the viral polymerase detaches from its template at a “break point” and resumes elongation at a downstream “rejoin point” by copying the 5’ end of the nascent daughter strand [12, 22], This process results in the formation of a new junction sequence and a truncated genome flanked by reverse complementary ends [23]. cbDVGs have long been thought to result from errors made by the RNA-dependent RNA polymerase (RdRps) during replication due to a combination of lack of proofreading activity and the presence of a polymerase with lower replication fidelity [12], No pattern or specific sequences for the break and rejoin points of cdDVGs have been reported so far.

Based on our consistent observations of discrete populations of cbDVG generated during RSV infections *in vitro* and *in vivo* [6], we set out to test the hypothesis that the generation of cbDVGs is not completely stochastic but instead is regulated by a carefully orchestrated process. Upon identification of cbDVG populations generated during infection, we show that specific viral sequences within the viral genome are preferred sites for cbDVG generation and that these sequences are conserved across viral strains. Utilizing this knowledge, we generated a recombinant virus that produced a restricted set of cbDVGs, serving as strong evidence that specific sequences dictate where cbDVGs are generated. In addition, we demonstrate for the first time that common cbDVGs are generated independently in natural infections in humans, further supporting an orchestrated origin for cbDVGs.

## Results

### High sensitivity detection of cbDVG populations during infection

To acquire a comprehensive view of the population of cbDVGs generated during infection, we developed an algorithm to identify cbDVGs junction regions within RNA-seq datasets with high sensitivity. The principle of this Viral Opensource DVG Key Algorithm (VODKA) is illustrated in Fig 1A. In brief, the break point (T’) and the rejoin point (W) are far apart in the parental viral genome, but in cbDVGs they become continuous and form the new cbDVG junction sequence when the viral polymerase (Vp) is released from the template and rejoins in the nascent strand. Since cbDVG junction sequences are absent in the full-length viral genome, VODKA identifies the sequence reads that capture junction sequences and then filters out false-positives reads that fully align to the reference genome.

**Fig 1.**
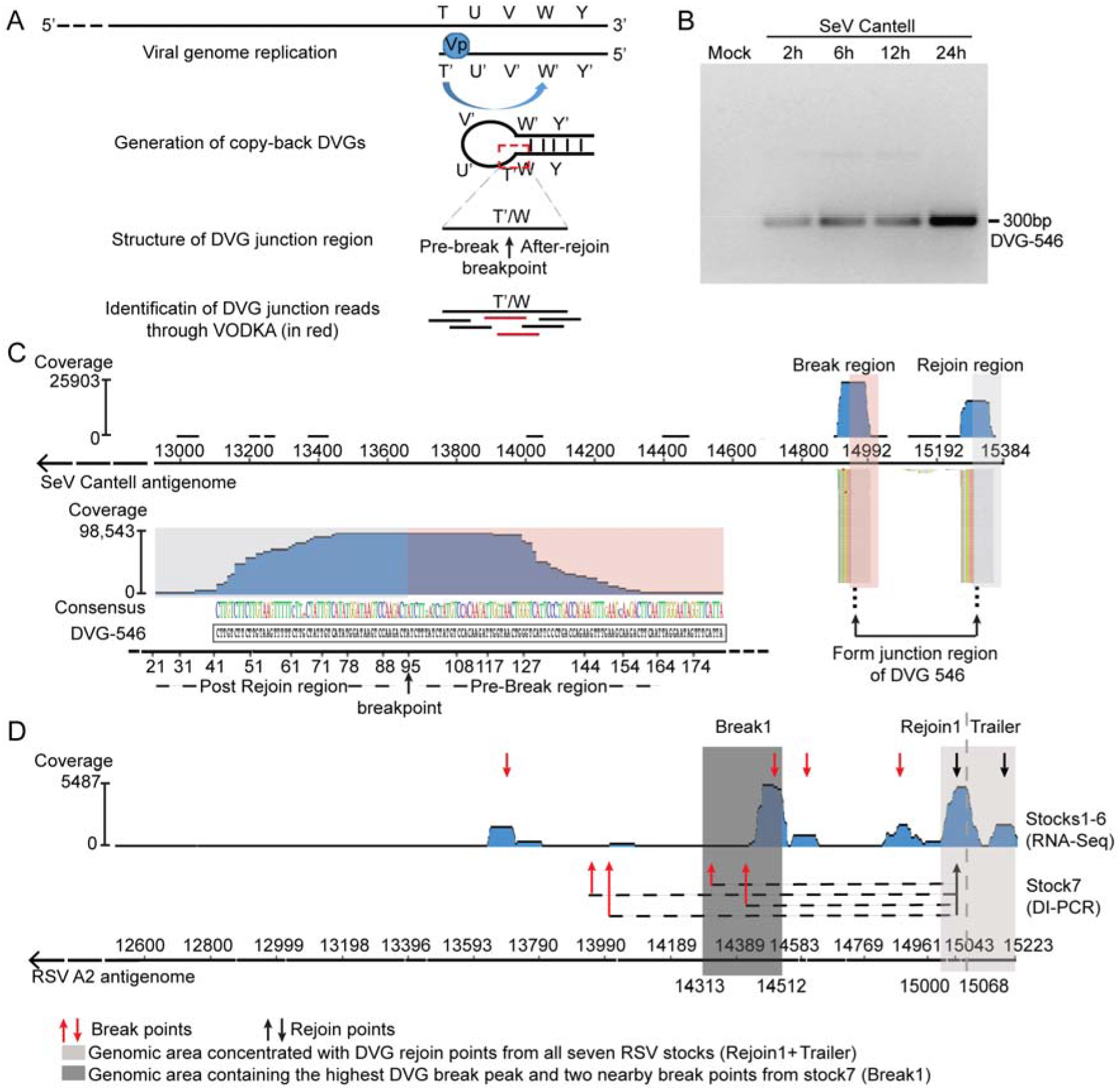
VODKA identified hotspots for polymerase break and rejoin during cbDVG formation. (A) Schematic representation of cbDVG generation from negative strand viruses and general principle behind VODKA identification ofcbDVGs. The red dashed square marks the unique junction region that distinguishes DVGs from full-length viral genomes. This junction region was used by VODKA to identify cbDVGs. (B) A549 cells were mock infected or infected with SeV Cantell at an MOI of 1.5 TCID_50_/cell and harvested at 2, 6, 12, and 24h post infection followed by detection of cbDVGs by RT-PCR using primers SeV Dll and gSeV Dll (SI Table). (C) Alignment of SeV cbDVG junction reads obtained from VODKA to the last 3kb nucleotides of the SeV antisense genome (top) or the DVG-546junction sequence (bottom). Blue histogram shows a synopsis of total coverage at any given position of the last 3Kb of the SeV reference genome (top) or the DVG-546 junction sequence (bottom). The numbers on the right-side axis of the graphs represent the total number of reads at the position with the highest coverage. Colored nucleotides beneath the blue histogram represent the consensus sequences of cbDVG junction identified by VODKA. The size of the letter represents the degree of conservation at the site and if there were mutations at a certain position the most representative nucleotide was listed on top. Black nucleotides beneath colored nucleotides indicate the sequence of the DVG-546junction region obtained by Sanger sequencing. The grey boxed areas in the graphs mark the region after the rejoin point and the pink boxed area the DVG region before the break point. (D) A549 cells were infected with RSV stocks1-7. For stocks1-6, RNA was extracted from the supernatants of the infected cells at 48 h post infections, followed by RNA-seq and VODKA screening. D VG junction reads were then mapped to the antigenome of the RSVA2 reference strain. Blue histogram shows synopsis of total coverage at any given position of the last 3kb of the RSV reference genome. The number on the right side of the graph represents the total reads at the position with highest coverage. Based on the VODKA output, individual peaks were identified as break or rejoin regions indicated by red or black facing-down arrows, respectively. For stock7, RNA was extracted from infected cells at 24 h post infections, followed by DVG specific RT-PCR using primer Dl1 and Dl-R (see S1 Table) that captures cbDVGs larger than 453 bp and with the break after Dl1 and the rejoin after Dl-R. Based on conventional cloning and Sanger sequencing ofPCR products, the break and rejoin points of cbDVGs detected in this stock were marked by red and black facing upwards arrows, respectively. Dashed lines indicate join sequences from individual cbDVGs in stock7. Light and dark grey shades represented the rejoin and break regions that were cloned into RSV minigenome backbone for further analysis in Fig 2A. Reference genome is shown at the bottom of the figure and the corresponding positions of DVG break/rejoin regions from all stocks, Break1, Rejoin1, and Trailer are also indicated.

We corroborated VODKA’s performance by testing for the presence of the highly dominant DVG-546 in samples from infections with SeV Cantell (Fig IB). Briefly, 98,543 out of the 98,626 (99.9%) cbDVG junction reads identified by VODKA from a RNA-seq data set obtained from SeV Cantell-infected cells mapped exactly to the known junction region of DVG-546 (Fig 1C, bottom panel). By aligning the cbDVG junction reads to the SeV full-length antisense genome, we determined the location of the major break and rejoin regions (blue peaks in the upper panel of Fig 1C). Each read that aligned to a break or rejoin region contained two portions, one of which fully aligned to the reference genome. The aligned reads in the break region (pink box, Fig 1C) and in the rejoin region (gray box, Fig 1C) corresponded to the DVG-546 sequence before the break point and after the rejoin point, respectively. The breakpoint for DVG-546 predicted by VODKA (14932±1_15292±1) exactly matched the one identified by Sanger sequencing (**↑** in Fig 1C), thereby establishing the efficiency and accuracy of VODKA in identifying cbDVG-specific sequences.

We then used VODKA to identify the population of cbDVGs generated during RSV infection. RSV is a virus known to generate immunostimulatory cbDVGs in infected patients [6], and thus a subject of interest in our laboratory. We analyzed pooled RNA-seq datasets from six RSV-infected cell cultures. These cultures were infected with RSV generated from the same parental stock (strain A2) that was first depleted of DVGs and then passaged independently in different cell lines to generate six different stocks enriched for DVGs. The presence of cbDVGs in these infections was confirmed using a specific RT-PCR followed by Sanger sequencing. By aligning VODKA-identified cbDVG junctions to the RSV A2 reference antigenome, we observed 4 major break hotspots spanning over 1300 nucleotides (red down-facing arrows in Fig ID). In contrast, only 2 major rejoin hotspots were observed within a narrower region of 223 nucleotides in length at the 3’end of the viral genome (black down-facing arrows in Fig ID). Remarkably, the rejoin area with the highest peak included counts present in all six virus stocks. We then compared these break and rejoin hotspots to those generated in infections with a different stock of RSV enriched in cbDVGs (stock7) from which the major cbDVGs were identified upon Sanger sequencing of PCR amplicons. We observed that the cbDVG rejoin points from stock7 were located within the strongest rejoin hotspot, whereas its four break points were distributed more broadly across the genome (Fig ID). These results reveal strong hotspots for the polymerase rejoin during cbDVG formation and suggest a large degree of conservation of the RSV cbDVGs rejoin positions.

### Identification of specific RSV genomic regions responsible for cbDVG generation

To determine if candidate hotspots were involved in cbDVG generation, we selected the region containing most break points (Break1, dark grey in Fig 1), or most rejoin points (Rejoin1+Trailer, light grey in Fig 1D) for further testing using a minigenome system [24], We constructed a RSV minigenome backbone (BKB) that included the reporter gene mKate2 for flow cytometry quantification of transcripts produced by the viral polymerase. In addition, the minigenome BKB included restriction enzyme sites to insert the selected Break1 and/or Rejoin1 regions (Fig 2A). The goal was to use this system to establish whether sequences in the candidate break and rejoin regions altered the polymerase elongation capacity, eventually leading to the generation of cbDVGs. As illustrated in Fig 2B, in this system mKate2 expression should only occur if the viral polymerase replicates the entire minigenome sequence from the trailer to the leader. Co-transfection of the minigenome construct with the four helper plasmids expressing the polymerase proteins (L, P, NP, and M2-1), resulted in mKate2 expression in 8-17% of the cells, whereas no mKate2 expression was detected in control transfections that lacked the viral polymerase (Fig 2C; -Vp). Constructs containing only the Rejoin1 sequence led to similar mKate2 expression as the BKB construct, whereas constructs containing Break1 caused a ∼30% reduction in mKate2 expression (Fig 2D-2E). We verified that the difference in mKate2 expression among transfections with different constructs was not due to variable transfection efficiency (S1A-S1C Fig), or cell death (S1D Fig). These results are consistent with the concept that during cbDVG generation, the viral polymerase falls off the template at the break region leading to a reduced amount of newly synthetized template available for mKate2 transcription.

**Fig 2.**
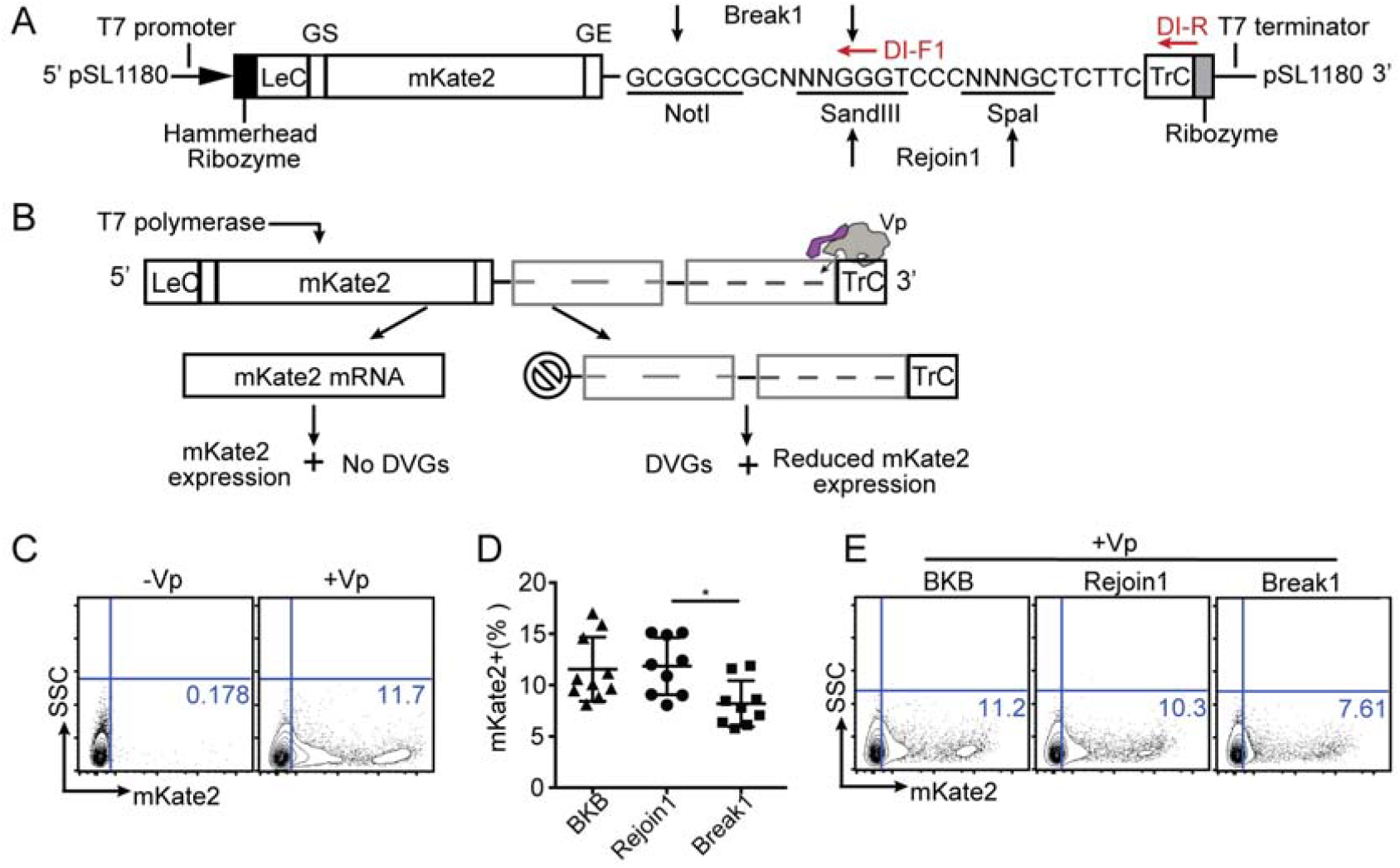
Specific RSV genomic regions alter the viral polymerase elongation capacity. (A) Schematic representation of the RSV minigenome backbone construct. Going from 5’ to 3’: T7 promoter, hammerhead ribozyme, complementary sense RSV leader sequence (LeC), NS1 gene start sequence, NS1 non-coding sequence, gene encoding mKate2 [25], L gene start sequence (GS), L gene end sequence (GE), restriction enzyme sites, complementary sense trailer sequence (TrC), hepatitis delta virus ribozyme, and T7 terminator. All constructs were generated in a pSH180 vector and included three restriction enzyme sites that were used for ligating Break1 and Rejoin1 as indicated. The length of all minigenome constructs after T7 promoter and before T7 terminator obeyed the rule of six. Red arrows indicate the primers that were used for detecting cbDVGs by PCR. One primer was within the viral trailer and the other primer was designed against the 3’ end of the break region. (B) Workflow of the RSV minigenome system. BSR-T7 cells were transfected with expression vectors encoding the polymerase genes M2-1, L, NP, P, and the minigenome construct to be tested. The minigenome was then transcribed by the T7 polymerase generating a template for amplification by the viral polymerase. If the paired junction region was not functional, the viral polymerase continued to elongate from 3’ TrC to the 5’ LeC, exposing the promoter in Lefor transcription leading to mKate2 expression. If break and rejoin sequences were functional, the viral polymerase recognized those signals resulting in dropping off from the template, generation of cbDVG-like fragments, and reduction of mKate2 expression. Since DVG junctions are split into two non-continuous regions (break and rejoin) in the minigenome plasmid, detection of the end-to-end DVG junction regions via DI-RT-PCR is a second indication of a productive junction region. (C) BSR-T7 cells were transfected with BKB either with or without the polymerase expression plasmids. One representative plot of mKate2 expression determined by flow cytometry is shown. (D-E) BSR-T7 cells were co-transfected with all 4 helper plasmids as well as either a BKB construct or a construct including the candidate signal sequences. mKate2 expression was measured by flow cytometry. The quantification of all repeats is shown as a percentage in D (F test P=0.0028). Representative flow plots are shown in E. *p<0.05 by one-way ANOVA Bonferroni’s multiple comparison test, n=8, mean±SEM, Bartlett’s test is used for examination of equal variance. Flow gating strategy is shown in S1A Fig.

To formally assess whether candidate break and rejoin sequences lead to cbDVG formation, we cloned the designated Pair1 composed of Break1/Rejoin1 into the minigenome system. Upon transfection, we observed that the construct containing Pair1 led to a similar degree of mKate2 expression than the construct bearing Break1 alone (Fig 3A-B). We also observed two major amplicons (white arrows in Fig 3C), both of which were absent in cells transfected with the construct bearing Break1 alone. These two amplicons contained cbDVGs that were confirmed by conventional Sanger sequencing (S2A Fig). The individual break and rejoin points of these minigenome-generated cbDVGs are indicated in Fig 3D. Interestingly, the rejoin points clustered in close proximity to the rejoin points that we identified from *in vitro* infected cells. Taken together, these data demonstrate that RSV cbDVG rejoin points fall into a discrete region of the viral genome, which is critical for cbDVG generation.

**Fig 3.**
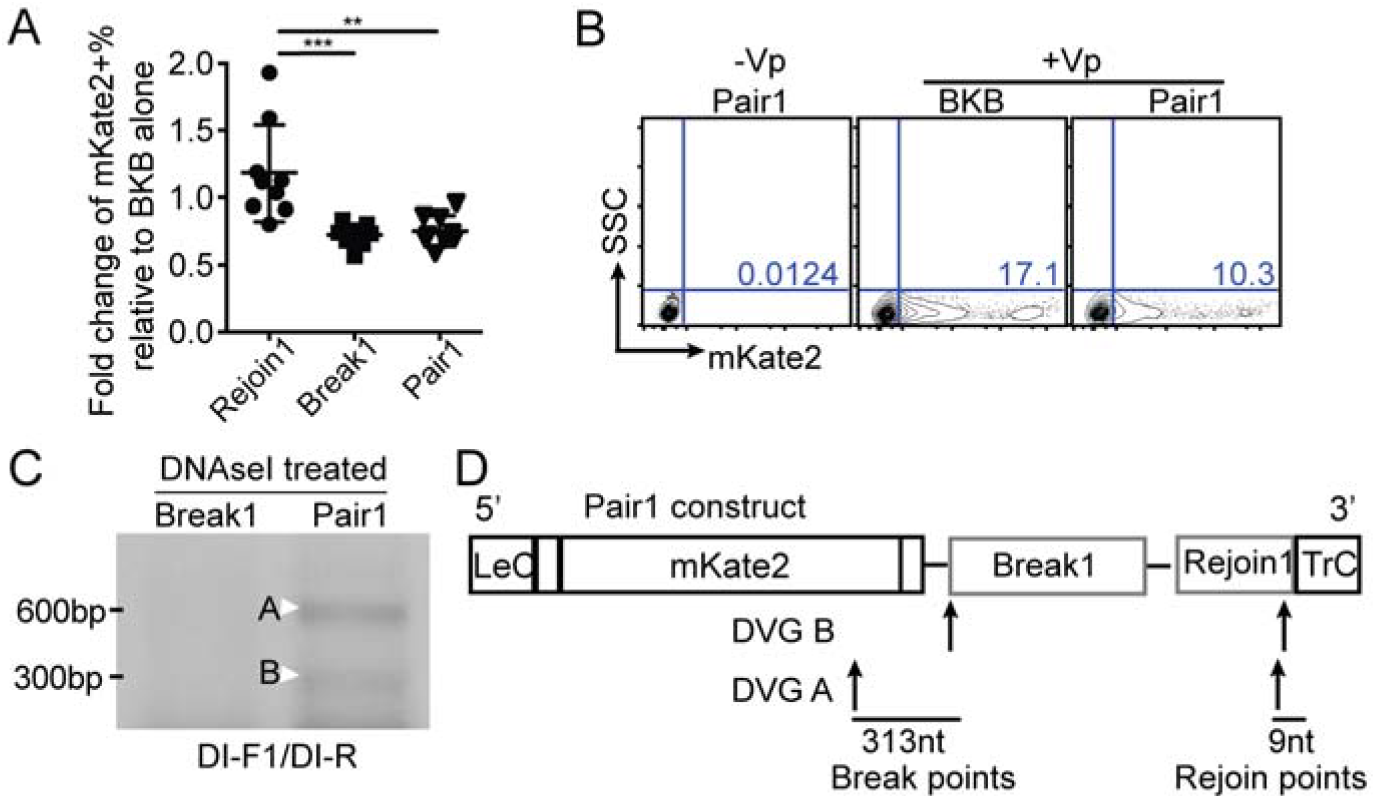
Identification of one conserved genomic region critical for cbDVG rejoin. (A-B) BSR-T7 cells were co-transfected with all **4** helper plasmids as well as BKB or constructs with candidate signal sequences. mKate2 expression was measured by flow cytometry. The quantification of all repeats is shown as a fold change expression over BKB in A (F test P<0.0001). **p<0.01, ***p<0.001 by one-way ANOVA Bonferroni’s multiple comparison test, n=8, mean±SEM, Bartlett’s test is used for examination of equal variance. Flow gating strategy is shown in S1A Fig. Representative flow plots are shown in B. (C-D) cbDVG-like fragments formed in Pair1 were detected by RT-PCR using DI-F1 and DI-R primer sets (S1 Table). Representative gel picture was shown in C. Bands labeled with a white arrow were confirmed to correspond to cbDVG by conventional sequencing. Their sequence is shown in S2A Fig. Location and the sequence length between the break and rejoin points are indicated in panel (D).

### Specific nucleotide composition determines the position for cbDVGs rejoin

Since the late Rejoin1 + early Trailer region of the RSV genome was highly enriched with DVG rejoin points relative to other regions in the RSV genome, we then examined which specific features within this region impacted cbDVG generation. This region is within one of most A and U enriched areas of the RSV genome (Fig 4A), suggesting that nucleotide composition might play a role in directing cbDVG formation. To avoid affecting the L gene “end signal” and the genome trailer region, we chose to mutate six nucleotides at the beginning of this rejoin region (nucleotide positions 191-186 from the 3’ end of antigenome) to either all Us (named GC>Us) or all GCs (named AU>GCs). We then used RT-PCR (DI-1/DI-R primer set) to detect cbDVG-like fragments formed in the cells co-transfected with all Us or all GCs mutant constructs and polymerase-expressing plasmids, as described earlier. Mutant GC>Us generated a dominant amplicon (lane2, Fig 4B) that was absent in cells transfected with mutant AU>GCs (lane4, Fig 4B). From sequencing PCR products within the areas marked by asterisks in Fig 4B, we identified five distinct rejoin points from mutant AU>GCs and three from mutant GC>Us (**↑** in Fig 4C). Compared to WT Pair1, the mutant GC>Us did not generate rejoin points proximal to the mutated region (grey area in Fig 4C), whereas the mutant AU>GCs still produced cbDVGs-like fragments at the mutated area.

**Fig 4.**
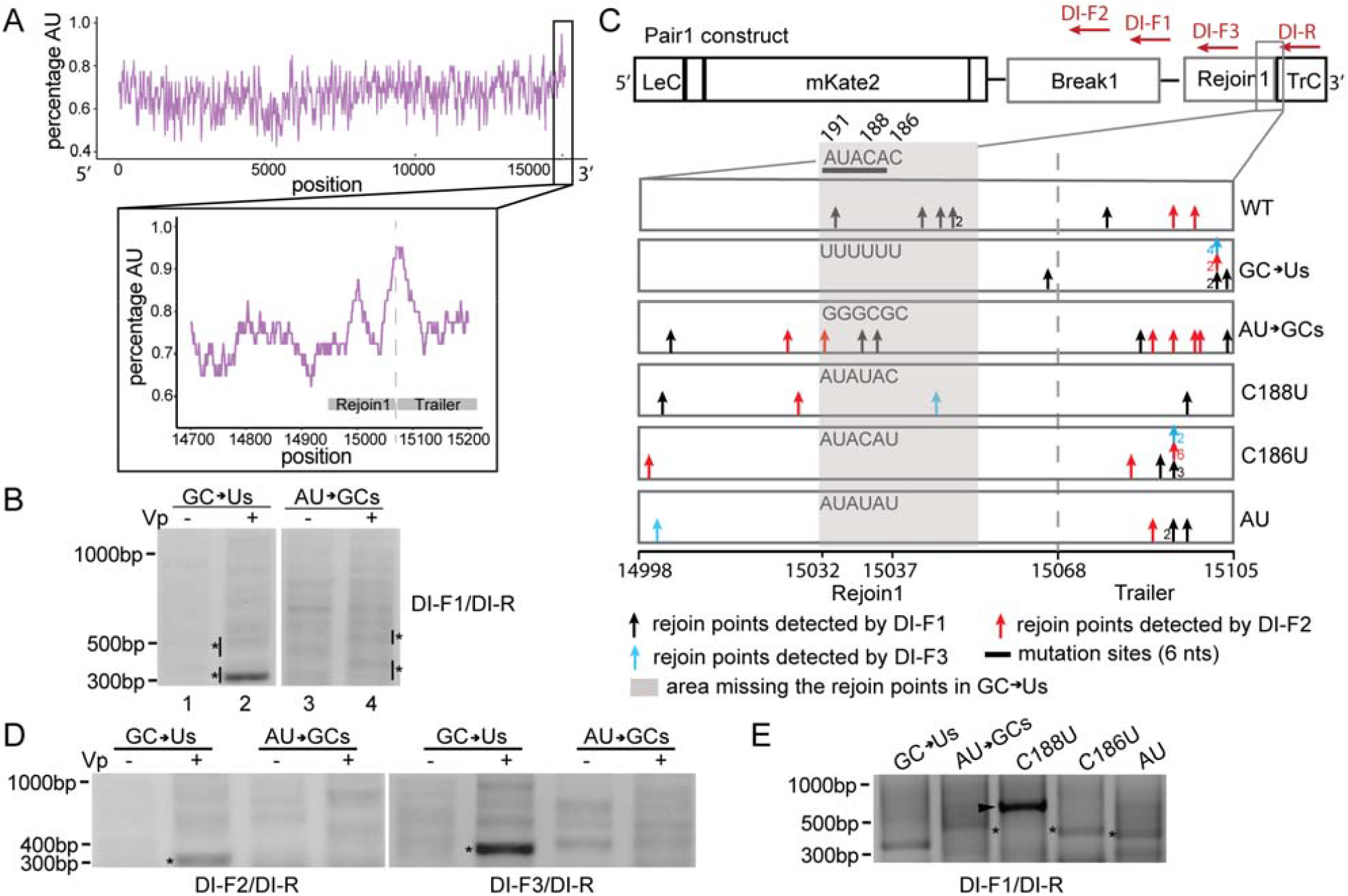
Specific nucleotide composition determines the position for cbDVGs rejoin. (A) AU-content (top, purple) over the entire genome and major rejoin region (inset, purple) of the RSV strain A2 antigenome. AU-content is plotted centered on a 40 base-pair sliding window in both plots. Blue ticks in the inset represent rejoin points from stock1-6 at each position starting at base 14700, with a proportionally darker color representing increased rejoin frequency at a given position. Grey box indicates the position ofRejoin1 and Trailer. (B) cbDVG-like fragments generated in mutant GC>Us and mutant AU>GCs were detected by RT-PCR using the DI-F1 and Dl-R primer set (S1 Table). Confirmed cbDVG-like fragments are indicated with asterisks next to the gel. (C) Schematic summary of all identified cbDVG rejoin points within positions 14998-15105 of the genome (RSVA2, reference genome NCBI KT992094.1) generated from WT Pair1 and Pair1 mutants minigenomes. This region includes gene end of L and the beginning of Trailer. Different colored arrows indicate rejoin points from cbDVGs detected by different primer sets as indicated in the figure. Primer sets are composed of the identical reverse primer, Dl-R, and one of either three different forward primers, DI-F1, DI-F2, and DI-F3. The numbers near the arrows indicated the number of times (>1) the same cbDVG was observed at this location (out of three experimental repeats with 1-3 clones sequenced each time). The thick black line marks the sequence of rejoin region selected for mutation studies. The antigenomic position range of the mutations is indicated at the bottom of the figure and the actual RNA sequences at the mutation sites in WT and five mutants are indicated. The numbers on top of the WT RNA sequence indicate their positions from the 3’ end of the antigenome. Grey shade marks the area where mutants GC>Us, C186U, and AU, did not have the rejoin points. (D) cbDVG fragments generated in mutant GC>Us and mutant AU>GCs were detected by DI-F2/DI-R and Dl-F3/DI-R through DVG specific RT-PCR. The DVG-like bands labeled by asterisks were confirmed to contain the same cbDVG-like fragment detected by DI-F1/DI-R in B (only band labeled by asterisk). (E) cbDVG fragments generated in mutants C188U, C186U, and AU were detected by DVG specific RT-PCR using the indicated primers. Confirmed cbDVG-like fragments are indicated with asterisks. Band labeled with an arrowhead was confirmed not to be a DVG product. Sequences of all the confirmed cbDVG-like fragments are shown in S2B-S2F Fig.

To rule out bias due to primer location, we designed two additional forward primers to detect cbDVG-like fragments from the same samples. Rejoins detected with DI-F2 primers are identified with red arrows in Fig 4C, while rejoins detected with DI-F3 are indicated with blue arrows. Transfections with mutant GC>Us led to one strong amplicon while not predominant amplicon was observed in transfections with mutant AU>GCs (Fig 4D), agreeing with results obtained using the DI-F1 primer set. Sequencing confirmed that the strong amplicons produced by all three different primer sets in transfections with mutant GC>Us were analogous cbDVG-like fragments and shared their break and rejoin points (sequence in S2 Fig, DVG 303bp). To examine if the observed lack of a predominant product resulting from mutant AU>GCs was due to a general reduction of replication ability of the viral polymerase induced by mutations, we introduced the same mutations in the construct with Rejoin1 alone and examined mKate2 expression by flow cytometry. We found no significant differences between Rejoin1 and the two mutants, Rejoin1-GC>Us and Rejoin1-AU>GCs. Neither of these constructs reduced mKate2 expression compared to BKB (S3 Fig), suggesting that the function of the RSV minigenome system remained intact despite of the mutations. Altogether, these data suggest that a minimal content of GC nucleotides in the rejoin region determines if cbDVGs are produced at that particular genomic location.

To determine if any of the two Cs within the mutated sequence was critical for cbDVG rejoining at this location, we performed a similar analysis using three new constructs: first C at position 188 or second C at position 186 from the 3’ end of antigenome mutated to U (named C188U or C186U, respectively), or both Cs mutated to Us (named AU). Transfection of the C186U, but not the C188U construct, resulted in one major DVG amplicon (indicated with an asterisk in Fig 4E; sequences in S2 Fig). The C186U construct rejoin points skipped the mutation area and concentrated in the early trailer region, similar to GC>Us. This was confirmed by the two other primer sets. A strong band shown in lane C188U at high molecular weight (indicated with an arrowhead in Fig 3E) was determined to not correspond to a cbDVG by Sanger sequencing. The construct bearing the double mutation (AU) behaved similar to C186U in terms of the rejoin positions (Fig 4C). Thus, we found the second C at position 15037 (position 186 from 3’ trailer end of antigenome) to be critical for cbDVG generation.

### A conserved rejoin region determines cbDVG formation during viral infection

To next establish whether rejoin1 impacts on cbDVG generation during viral infection, we created a mutant virus harboring mutations identical to the GOUs minigenome construct. This virus is herein identified as gRSV-FR-GC>Us. The backbone of the recombinant RSV (Line 19) included the mKate2 gene and we used mKate2 expression to estimate its replication. As shown in Fig 5A, cells infected with gRSV-FR-GC>Us expressed the same level of mKate2 protein as cells infected with the WT reporter virus (gRSV-FR-WT) at 72 h post infection. Both viruses began to generate cbDVGs from P3 and the pattern of cbDVGs was maintained, and became stronger, in P5 (Fig 5B). We verified that P5 gRSV-FR-GC>Us still carried the mutations we introduced (Fig 5C). Interestingly, gRSV-FR-WT produced 4 major DVGs, whereas gRSV-FR-GC>Us only generated one dominant cbDVG (asterisks in Fig 5B, confirmed sequence in S2G and S3H Fig), which is consistent with results from the minigenome system. The dominant cbDVG generated in cells infected with gRSV-FR-GC>Us rejoined at the early trailer region and skipped the mutation site, similar to what was observed in the minigenome system. Cells infected with gRSV-FR-WT produced one cbDVGs that rejoined within the mutation site and other 3 cbDVGs that rejoined at the same region of the mutant virus (Fig 4D).

**Fig 5.**
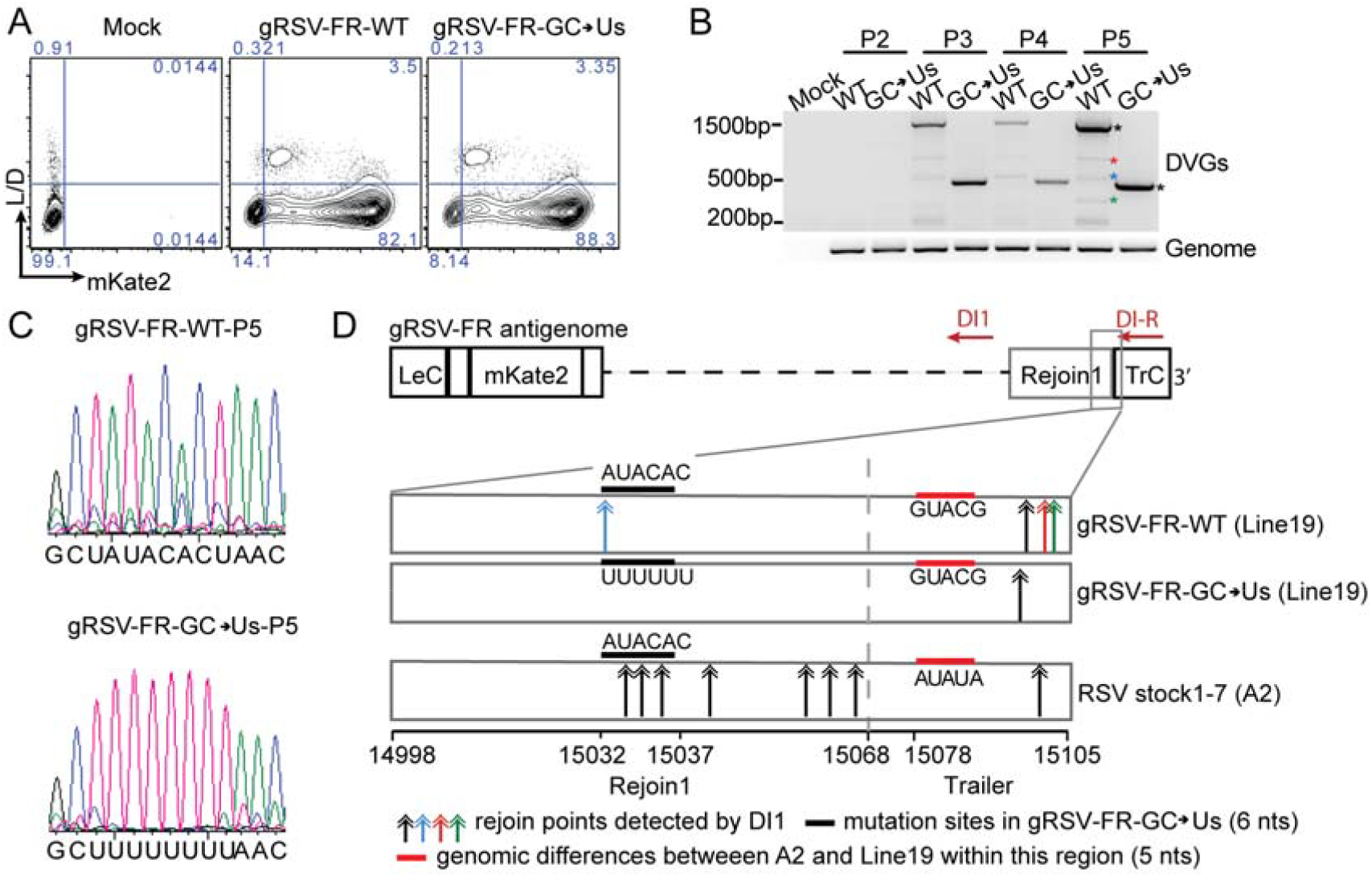
A conserved rejoin region determines cbDVG formation during viral infection in tissue culture. (A) HEp2 cells were infected with gRSV-FR-WT-P3 and gRSV-FR-GC->Us-P3 at MOI of 1.5 and harvested at 72 h post infection and stained for aqua L/D, followed by flow cytometry for mKate2 expression as a proxy for virus replication. (B) cbDVG detection from P2, P3, P4, and P5 of gRSV-FR-WT and gRSV-FR-GC>Us via DVG specific RT-PCR using DI1/DI-R primer set in HEp2 cell. Confirmed cbDVG-like fragments are labeled by asterisks next to the gel. Different colors correspond to the color of double arrows in Fig 4D. Sequence of confirmed DVGs are listed in S2G and S2H Fig. (C) Confirmation of introduced mutations in gRSV-FR-GC->Us-P5 infected cells via Sanger sequencing. (D) Schematic summary of all identified cbDVG rejoin points within positions 14998-15105 (RSVA2, reference genome NCBI KT992094.1) generated during infections with gRSV-FR-WT, gRSV-FR-GC>Us, and RSV stocks1-7. All DVGs were detected by DVG specific RT-PCR using the DI1/DI-R primer set. RSV HD stock1-7 are strain A2 and gRSV-FR-WT and gRSV-FR-CG->Us are strain Line19. The sequence difference between A2 and line 19 within common rejoin region is indicated by a red line. Colored double arrows indicated the rejoin points of cbDVGs detected from viruses. Black parallel line marks the 6 nucleotides mutated in gRSV-FR-GC->Us.

The majority of rejoin points found in infection with gRSV-FR-WT, which derived from RSV Line 19, located within the early trailer sequence, rather than around the mutation site as found in infection with RSVstocksl-7 derived from RSV line A2 (Fig 5D). Alignment of both sequences revealed one natural mutation in RSV Line 19 that introduced three GCs right at the beginning of the trailer sequence, which are not present in RSV A2 (sequence indicated with a red horizontal line in Fig 5D). The increased GC content in this position in Line 19 likely explains why gRSV-FR-WT generates more cbDVGs at this location than RSV A2 stocksl-7. Regardless of this natural preference for rejoining in the early trailer, gRSV-FR-GC>Us diminished the rejoin signal at the mutation site as no cbDVGs rejoin points were found at this location and resulted in less diversity of cbDVG generation compared to the WT virus. Overall, these data confirm that the common rejoin region sequence tested in the minigenome system determines cbDVG rejoining during RSV infection and that the GC content on this region critically determines the site of cbDVG rejoin.

### Conserved cbDVGs form during natural RSV infections in humans

To examine whether the rejoin1 region was utilized during natural infections, we applied VODKA to RNA-seq datasets obtained from RSV-positive pediatric clinical samples. A total of 10 pediatric specimens were sequenced; 4 were classified as DVG-low and 6 as DVG-high based on semi-quantification following cbDVG PCR. VODKA outputs were aligned to the reference genome of an RSV strain A isolate (Reference genome NCBIKJ672447, 2012) and showed that, consistent with previous cbDVG-RT-PCR results, DVG-low samples (upper panel in Fig 6A) contained ∼8 fold less cbDVG junction reads than DVG-high patients (lower panel in Fig 6A). In addition, the coverage mapping showed the presence of multiple break and rejoin regions. Some of them were a mix of both break points and rejoin points (Fig 6A, read and black arrows). The rejoin points were particularly noteworthy because the majority of them clustered within one narrow AU-rich “Rejoin1+ Trailer” region (red ticks in Fig 6B) similar to that identified in *in vitro* infections (blue ticks in Fig 6B). According to the frequency of different cbDVG junction positions, we illustrated the top 6 major cbDVGs (one of them is a snap-back) in Fig 6C (details summarized in Table 1, Break_Rejoin position shown as T’_W). All of them were found in multiple patients (Fig. 6C and Table 1). The most abundant DVG again had the rejoin point within the “Rejoin1+Trailer” region, despite of a higher diversity of rejoin points compared to *in vitro* infection, possibly due to the deeper depth of RNA-seq used to analyze clinical samples. Taken together, these results demonstrate that a conserved rejoin region drives the generation of most cbDVGs during RSV infection *in vitro* and *in vivo* and that identical RSV cbDVGs are generated in naturally infection of different individuals

**Table 1.**
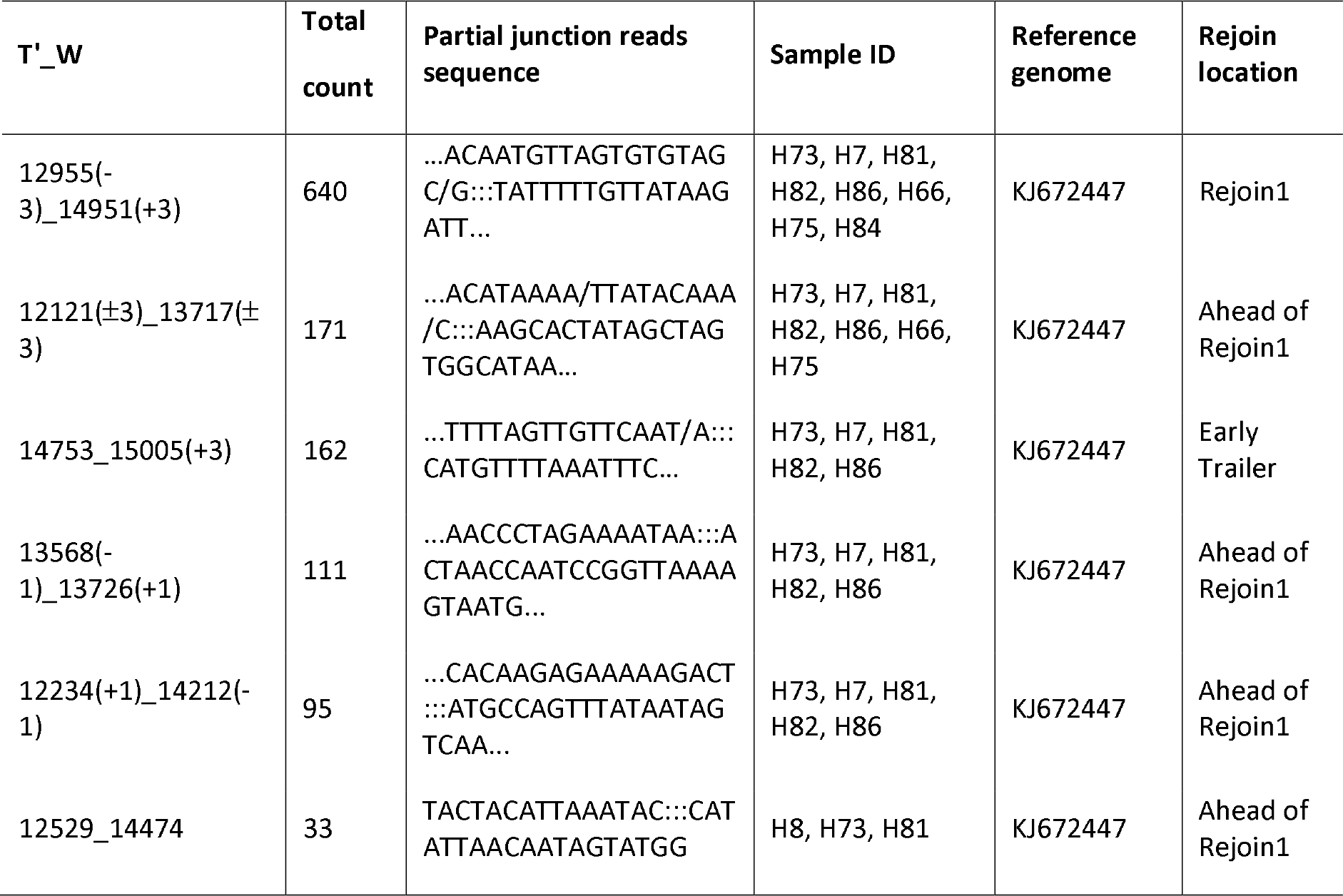
Major DVG junctions identified by VODKA from RSV positive nasal secretions.

**Fig 6.**
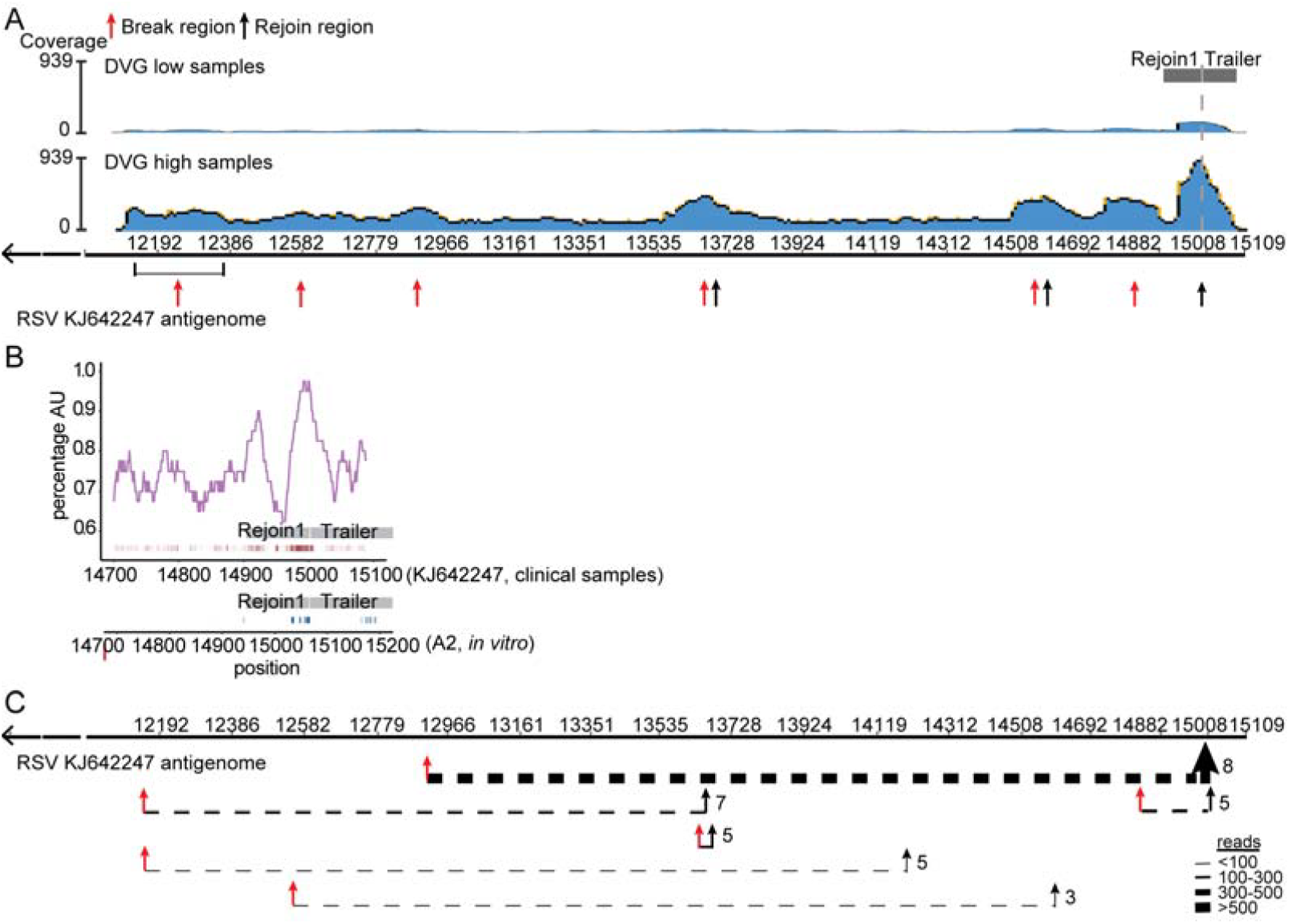
A conserved rejoin region determines cbDVG formation in infected human patients. (A) Four specimens from RSV A-positive pediatric patients with low contents of DVGs and 6 with high content of DVGs were selected for RNA-seq. All seq reads that passed the quality threshold were analyzed by VODKA to screen for cbDVG junctions. DVG junction reads were then aligned to the RSV reference genome NCBIKJ672447. Blue histogram shows a synopsis of total coverage at any given position of the RSV reference genome (illustrated by the y-axis). The number on the right side of the graph represents the total reads at the position with highest coverage. (B) AU-content (top, purple) within nucleotides 14700-15109 of the RSV strain A isolate KJ642247 (human samples) is plotted centered on a 40 base-pair sliding window in both plots. Grey box indicated the position of Rejoin1 and Trailer. Individual rejoin points identified by DVG junction reads from pediatric samples within this region are represented by red ticks underneath the graph. The rejoin points from in vitro infection of RSV stocks 1-6 are plotted using blue ticks. The darker color represented increased rejoin frequency at a given position. (C) Top 6 major DVGs identified from clinical samples are depicted by dashed lines that join their break points (red arrows) and rejoin points (black arrows). The thickness of the line represents the frequency of DVG reads at a given DVG as indicated. The number of patients that have the Particular DVG is indicated at the right of the rejoin point for each cbDVG.

## Discussion

DVGs are critical regulators of viral replication and pathogenesis in multiple RNA virus infections, but the mechanisms modulating their generation are unknown. Historically, DVGs are thought to result from random errors introduced during replication by the viral polymerase. However, mounting evidence indicates that the generation of cbDVGs is not totally stochastic. Specifically, we show that during RSV infection discrete hotspots in the viral genome mark sites of the viral polymerase release and rejoin during cbDVG formation, both *in vitro* and during natural RSV infections in humans. Moreover, we show that the content of GC nucleotides within the major rejoin hotspot critically impacts the generation of cbDVGs at that position and we identified specific nucleotides that, when mutated, altered the ability of recombinant viruses to generate diverse species of DVGs. The identification of a specific sequence involved in cbDVG formation opens the unprecedented possibility of genetically manipulating the content of cbDVGs during infection. This possibility may significantly impact our ability to generate tools to further understand the role of these viral products in virus pathogenesis, as well as potentially manipulate the cbDVG content with antiviral and/or therapeutic purposes.

In this study, we utilized a custom-designed algorithm, VODKA, to identify cbDVG in infections *in vitro* or *in vivo* from children naturally infected with RSV. VODKA outputs were consistent with previous results obtained using classic DVG-RT-PCR and demonstrated a higher sensitivity in the detection of cbDVGs both *in vitro* and in clinical samples. False-positive DVG junction reads were ruled out by screening all reads aligned to the host (reads from human transcriptome) using VODKA. This test resulted in a minimal number of hits compared to viral samples, adding to the evidence reported throughout this manuscript to support the specificity of cbDVG detection by VODKA. VODKA can successfully identify cbDVGs in a number of viruses, including SeV (Fig 1) and Ebola (not shown), offering a powerful tool for cbDVG detection in clinical samples.

Based on our data, we conclude that the rejoin position significantly influences cbDVG generation. A minimum proportion of GC nucleotides are necessary for rejoining to occur at that particular region, and one nucleotide alone can influence the location of the DVG rejoin point implying that a strong rejoin signal likely needs an optimal number of GC nucleotides in specific locations. Importantly, the same differential distribution of cbDVG rejoin points was observed when we compared cbDVG generation from infections with RSV A2 and Line19, which differ in their GC content at the beginning of the trailer region. In addition, our data suggest that rejoin sequences influence the function of break signals when inserted as pairs in the construct. Our data is in agreement with data from *in vitro* infections with measles virus lacking the C protein, where break points of cbDVGs were widely distributed along the genome, whereas the rejoin points were clustered in a narrow region close to 5’ end of the genome [26].

Further investigation into the molecular details of how the viral polymerase recognizes these signals may lead to important insights about the mechanism involved in RSV virus replication and the generation of cbDVGs. A lower density of nucleocapsid proteins (NPs) at certain genomic locations has been shown to result in increased cbDVG formation in SeV infection [27], However, the mutations described to be responsible for low NP density were absent in our SeV stocks, suggesting that alternative mechanisms are likely involved. The usage of C nucleotides as a signal closely resembles the recognitions of “gene end” or “gene start” by the viral polymerase when working on transcription [28, 29] and it would be intriguing to evaluate if the mechanisms of cbDVG generation and viral RNA transcription are related. Another factor influencing DVG accumulation is their length, which is tightly related to the spatial structure of the viral RNPs. In paramyxoviruses, although it is thought that “only genomes with hexametric or heptametric lengths are efficiently replicated” [30, 31], some cbDVGs generated *in vitro* do not obey this rule [26, 32, 33]. For RSV, we observed that a number of cbDVGs do not follow the rule of six or the rule of seven. Nonetheless, cbDVGs with certain length may have increased replication efficiency and thus an enhanced fitness advantage. Interestingly, in our minigenome system, although cbDVGs from Pair1 contained the expected rejoin point positions, break points frequently fell into a region further ahead of Break1, suggesting that the distance between the Break and Rejoin points may also play a role in determining where the break position is.

In addition to genomic sequences, other factors, such as viral proteins, likely play an important role in DVG generation. For instance, influenza viruses harboring a high fidelity polymerase generate fewer deletion DVGs [34], Mutations in non-structural protein 2 of influenza have also been shown to increase the *de novo* generation of DVGs by altering the fidelity of viral polymerase [35]. Host factors may be essential contributors to DVG generation as well [10]. For example, vesicular stomatitis virus produces a large amount of snap-back DVGs in most cell lines, except human-mouse somatic cell hybrids, and this cellular attribute was mapped to human chromosome 16 [36]. Similarly, infection with measles virus did not show *de novo* generation of DIPs in human WI-38 cells and SeV did not produce cbDVGs in chicken embryo lung cells [37, 38]. Despite the potential importance of these additional factors on DVG generation, the current work represents a major paradigm shift with the identification of sequences that regulate cbDVG formation.

Remarkably, we found various common cbDVGs present in more one than one patient and at least one of those cbDVGs was also present in infections *in vitro.* These observations support a conserved origin for cbDVGs during infection and challenge the idea that DVGs occur as random product of virus replication. To date, all studies on DVG biology have been correlative in nature. This work opens up new areas of investigation and can ultimately allow us to manipulate the ability of viruses to produce DVGs as a powerful tool to study the role of DVGs in viral pathogenesis.

## Materials and methods

### Ethics statement

Studies of human samples were approved by University of Pennsylvania Institutional Review Board. The embryonated chicken eggs used in theses studies were 10 days old and were obtained from Charles River.

### Cells and viruses

A549 cells (human type II alveolar cells, ATCC, #CRM-CCL185) and HEp2 cells (HeLa-derived human epithelial cells, ATCC, CCL23) were cultured at 7% CO_2_ and 37°C with Dulbecco’s modified Eagle’s medium supplemented with 10% fetal bovine serum (FBS), 1 mM sodium pyruvate, 2 mM L-Glutamine, and 50 mg/ml gentamicin. BSR-T7 cells (Hamster kidney cells, BHK cells constitutively expressing the T7 polymerase, provided by Dr. Christopher Basler’s lab at Icahn School of Medicine) and were maintained in 10% FBS with 1 mg/ml Geneticin (Invitrogen). All cell lines were treated with mycoplasma removal agent (MP Biomedicals) and routinely tested for mycoplasma before use. Sendai virus Cantell stock (referred to as SeV HD, with high DVG particle content) was prepared in embryonated chicken eggs as described previously [7, 39]. The SeV HD stock used in these experiments had a high infectious to total particle ratio of 500:15,000. RSV-HD stocks 1-7 (stock of RSV derived from strain A2, ATCC, #VR-1540 with a high content of cbDVGs) were prepared and characterized as described previously [6, 40] in MAVS KO, STAT1 KO, and WT A549 cells, respectively. The cell lines were kindly provided by Dr. Susan Weiss (University of Pennsylvania).

### Plasmids

Mammalian expression vectors for RSV N (NR-36462), P (NR-36463), M2-1 (NR-36464), and L (NR-36461) proteins, and the RSV reverse genetic backbone pSynkRSV-line19F (rRSV-FR, NR-36460) were obtained from BEI Resources. Detailed information of the constructs can be found in reference [41], The backbone plasmid of the RSV minigenome used for testing various DVG junction regions was constructed by cloning two regions of sequences amplified from pSynkRSV-line19F into the pSH180 vector. The first region included a T7 promoter, a hammerhead ribozyme, RSV leader sequence, and genes encoding monomeric Katushka 2 (mKate2), while the second region included the RSV trailer sequence, a Hepatitis delta virus ribozyme and a T7 terminator. These regions were sequentially cloned into pslll80 vector using restriction enzyme pairs Spel/Sandl and Sandl/EcoRI, respectively. The potential cbDVG break and/or rejoin regions (positions in S1 Table) were then inserted between those two regions using restriction enzyme pairs Notl/Sandl and Sandl/Spal, respectively. A detailed scheme of the construct can be seen in Fig 2A. Pair1 and Rejoin1 mutations were introduced using the site-directed mutagenesis commercial kit QuickChange® II XL (Agilent, CA) according to the manufacture’s protocol. All primers used for cloning are listed in S1 Table. Mutations in reverse genetic backbone pSynkRSV-line19F were generated by fusion PCR using primers in S1 Table as previously described [42],

### RNA extraction and DVG-RT-PCR

Total RNA was extracted using TRIzol or TRIzol LS (Invitrogen) according to the manufacturer’s specifications. For detection of RSV DVGs in RSV infection, 1-2 μg of isolated total RNA was reverse transcribed with the DI1 primer using the SuperScript III reverse transcriptase (Invitrogen) without RNase H activity to avoid self-priming. Recombinant RNase H (Invitrogen) was later added to the reverse transcribed samples and incubated for 20 min at 37°C. DVGs were partially amplified using both DI1 primer and Dl-R primer. The temperature cycle parameters used for the cbDVG-PCR in a BioRad C1000 Thermal Cycler were: 95°C for 10 min and 33-35 cycles of 95°C for 30 sec, 55°C for 30 sec and 72°C for 90 sec followed by a hold at 72°C for 5 min. Ramp rate of all steps was 3 degree/sec. Detailed method can be found in[6]. For detection of cbDVGs in the RSV minigenome system, extracted RNAs were treated with 2 μ1 TurboDNasel (Invitrogen) for 15 min at 37°C, followed by reverse transcription. Same procedures as above were utilized, except replacing DI1 primer with DI-F1, DI-F2, and DI-F3 primers. These were then all paired with DI-R reverse primer to amplify the different sizes of PCR products. Sequences of all primers are listed in S1 Table.

### RT-qPCR

Total RNA (1 μ.g) was reversed transcribed using the high capacity RNA to cDNA kit from Applied Biosystems. cDNA was diluted to a concentration of 10 μg/μl and amplified with specific primers in the presence of SYBR green (Applied Biosystems). qPCR reactions were performed in triplicate using specific primers and the Power SYBR Green PCR Master Mixture (Applied Biosystems) in a Viia7 Applied Biosystems Light-cycler. Gene expression levels of RSV G were normalized to the *GAPDH* copy number. Sequences of primers used in these studies can be found in S1 Table.

### RNA-Seq

RNA-Seq for SeV Cantell and RSV HD stocks 1-6 were performed as previously described [43]. RNA was extracted using TRIzol reagent and was re-purified using the PicoPure™ RNA isolation kit (Thermo Fisher Scientific). RNA quality was assessed using the RNA Pico 6000 module on an Agilent Tapestation 2100 (Agilent Technologies) prior to cDNA library preparation. For SeV RNA-Seq dataset, total cDNA libraries were prepared starting from 75 ng (SeV Cantell) and 450 ng (RSV HD stocks) of extracted raw RNA using the Illumina TruSeq Stranded Total RNA LT kit with Ribo-Zero Gold, according to the manufacturer’s instructions. Samples were run on Illumina NextSeq 500 to generate 75 bp, single-end reads, resulting in 21-53 million reads per sample, with an average Q30 score ≥ 96.8%. For sequencing of samples from RSV-positive patients, including 4 DVG low patients and 6 DVG high patients, 100-450 ng of extracted raw RNA was used for preparation of cDNA library using the same kit as above. Samples were run on Illumina NextSeq 500 to generate 150bp, paired-end reads, resulting in 60-170 million reads/sample with average Q30 score ≥ 84.6%. To analyze genomic AU-content relative to DVG break and rejoin points, we calculated the percentage of A or U nucleotides over sliding windows of 40 bases using the Python programming language (Python Software Foundation, https://www.python.org/). We plotted AU-content and cbDVG rejoin points in R using the ggplot2 package [44],

### Viral Opensource DVG Key Algorithm (VODKA)

Based on our *in vitro* RSV experiments, we made the assumption that most cbDVGs are generated from the viral sequence near the 5’ end region of the genome (close to the Trailer sequence). Therefore, starting with the last 3kb of a reference viral genome, we built an index of potential DVG sequences by taking all possible combinations of two non-overlapping segments of L bases, where L is the read length. The segments are linked by reverse complementing the second segment (C-D) and adding the first segment (A-B) to it (S4 Fig). Sequenced reads are aligned to the potential DVGs using bowtie2 [45], and subsequently undergo two filtering steps. First, reads are removed unless they map across a breakpoint (A_C) with at least 15bp of mapped segment on each side. Second, the reads that map cleanly to the reference genome are filtered out. This pipeline gives the output read counts for each breakpoint (A_C). To be consistent with the structure of copy-back DVGs in Fig 1A, C is equivalent to breakpoint T’ and A is equivalent to rejoin point W. VODKA output reads were further aligned to reference viral genomes (RSV A2: NCBI accession number KT992094.1; RSV 2012 clinical isolate: NCBI accession number KJ672447) or known SeV DVG-546 to identify the potential DVG junction regions using the Genious 7.0 software.

### RSV minigenome and reverse genetics system

BHK cells constitutively expressing the T7 polymerase (BSR-T7 cells) were transfected with different minigenome constructs, gRSV-FR-WT, or gRSV-FR-GC->Us, as well as the sequence-optimized helper plasmids encoding N, P, M2-1, and L, all underT7 control as described previously [41]. Cells were incubated with transfection complex (total plasmid: lipofectamine=1:3.3) for 2 h at room temperature and then at 37°C for overnight using Opti-MEM as medium. The following morning, the medium was replaced with antibiotic free tissue culture medium containing 2% FBS. For minigenome experiments, cells were harvested at 48 h post-transfection for either RNA extraction or flow cytometry. For mutant virus production, cells were maintained and split every 2-3 days until see obvious CPEs. Then viruses were collected and blindly passaged in HEp2 cells three times to obtain P3. P3 was titrated and passaged two more times at MOI of 10 to generate P4 and P5.

### Flow Cytometry

Transfected BSR-T7 cells were trypsinized 48 h post transfection and were either directly diluted in FACS buffer (PBS containing 20 mM EDTA) or stained with aqua L/D. Cells were washed twice in FACS buffer before flow cytometry analysis on an LSRFortessa (Becton Dickinson). Data analysis was performed using Flowjo version Legacy.

### Statistical analysis

All statistical analyses were performed with GraphPad Prism version 5.0 (GraphPad Software, San Diego, CA) and *R v3.4.1.* A statistically significant difference was defined as a *p*-value <0.05 by one-way analysis of variance (ANOVA) with a *post hoc* test to correct for multiple comparisons (based on specific data sets as indicated in each figure legend).

### Code availability

The VODKA algorithm is open-source and available at: https://github.com/itmat/VODKA.

### Data availability

All data are available upon request to the corresponding author. Raw RNA-Sequencing data of FISH-FACS sorted SeV infected cells and RSV infected samples have been deposited on the Gene Expression Omnibus (GEO) database for public access (SeV: GSE96774; RSV: GSE114948).

## Acknowledgements

This work was supported by the US National Institutes of Health National Institute of Allergy and Infectious Diseases (NIH AI083284 and AI137062 to C.B.L). We thank Dan Beiting for the support with RNA-seq and GEO submission.

Conceived experiments: Y.S., and C.B.L.; Developed methodology: Y.S., E.J.K, D.A., and G.R.G; Performed experiments, collected and analyzed data: Y.S., E.J.K., L.J.T., and S.F.; Provided reagents and resources: G.R.G; Wrote the original draft: Y.S., D.A., and C.B.L.; Supervised research activities: C.B.L

## Supporting information captions

S1 Fig. Identification of RSV genomic regions responsible for the generation of cbDVGs.

S2 Fig. Sequences of cbDVGs identified from labeled bands in RSV minigenome system and viral infection.

S3 Fig. GC mutations in the RSV minigenome did not alter viral polymerase processivity.

S4 Fig. Working scheme of VODKA.

S1 Table. Oligo primer list.

## References

1. Henle W, Henle G. INTERFERENCE OF INACTIVE VIRUS WITH THE PROPAGATION OF VIRUS OF INFLUENZA. Science. 1943;98(2534):87–9. Epub 1943/07/23. doi:10.1126/science.98.2534.87. PubMed PMID:17749157.

2. von MP. Propagation of the PR8 strain of influenza A virus in chick embryos. II. The formation of incomplete virus following inoculation of large doses of seed virus. Acta Pathol Microbiol Scand. 1951;28(3):278–93. PubMed PMID: 14856732.

3. Von Magnus P. Incomplete forms of influenza virus. Adv Virus Res. 1954;2:59–79. PubMed PMID:13228257.

4. Lopez CB. Defective viral genomes: critical danger signals of viral infections. J Virol. 2014;88(16):8720–3. doi:10.1128/JVI.00707-14. PubMed PMID:24872580; PubMed Central PMCID:PMCPMC4136278.

5. Weber M, Weber F. RIG-I-like receptors and negative-strand RNA viruses: RLRly bird catches some worms. Cytokine Growth Factor Rev. 2014;25(5):621–8. doi:10.1016/j.cytogfr.2014.05.004. PubMed PMID:24894317.

6. Sun Y, Jain D, Koziol-White cj, Genoyer E, Gilbert M, Tapia K, et al. Immunostimulatory Defective Viral Genomes from Respiratory Syncytial Virus Promote a Strong Innate Antiviral Response during Infection in Mice and Humans. PLoS Pathog. 2015;11(9):e1005122. doi:10.1371/journal.ppat.1005122.

7. Tapia K, Kim WK, Sun Y, Mercado-Lopez X, Dunay E, Wise M, et al. Defective viral genomes arising in vivo provide critical danger signals for the triggering of lung antiviral immunity. PLoS Pathog. 2013;9(10):e1003703. doi:10.1371/journal.ppat.1003703. PubMed PMID:24204261; PubMed Central PMCID:PMCPMC3814336.

8. Vasilijevic J, Zamarreno N, Oliveros JC, Rodriguez-Frandsen A, Gomez G, Rodriguez G, et al. Reduced accumulation of defective viral genomes contributes to severe outcome in influenza virus infected patients. PLoS Pathog. 2017;13(10):e1006650. Epub 2017/10/13. doi:10.1371/journal.ppat.1006650. PubMed PMID:29023600; PubMed Central PMCID:PMCPmc5638565.

9. Strahle L, Garcin D, Kolakofsky D. Sendai virus defective-interfering genomes and the activation of interferon-beta. Virology. 2006;351(1):101–11. doi:10.1016/j.virol.2006.03.022. PubMed PMID:16631220.

10. Frensing T. Defective interfering viruses and their impact on vaccines and viral vectors. Biotechnol J. 2015;10(5):681–9. doi:10.1002/biot.201400429. PubMed PMID:25728309.

11. Huang AS, Baltimore D. Defective viral particles and viral disease processes. Nature. 1970;226(5243):325–7. PubMed PMID:5439728.

12. Lazzarini RA, Keene JD, Schubert M. The origins of defective interfering particles of the negative-strand RNA viruses. Cell. 1981;26(2 Pt 2):145–54. PubMed PMID:7037195.

13. Pathak KB, Nagy PD. Defective Interfering RNAs: Foes of Viruses and Friends of Virologists. Viruses. 2009;1(3):895–919. doi:10.3390/v10308951. PubMed PMID:21994575; PubMed Central PMCID:PMCPMC3185524.

14. Janda JM, Davis AR, Nayak DP, De BK. Diversity and generation of defective interfering influenza virus particles. Virology. 1979;95(1):48–58. Epub 1979/05/01. PubMed PMID:442544.

15. Nayak DP, Chambers TM, Akkina RK. Defective-interfering (DI) RNAs of influenza viruses: origin, structure, expression, and interference. Curr Top Microbiol Immunol. 1985;114:103–51. PubMed PMID:3888540.

16. Kajigaya S, Arakawa H, Kuge S, Koi T, Imura N, Nomoto A. Isolation and characterization of defective-interfering particles of poliovirus Sabin 1 strain. Virology. 1985;142(2):307–16. Epub 1985/04/30. PubMed PMID:2997988.

17. Poirier EZ, Mounce BC, Rozen-Gagnon K, Hooikaas PJ, Stapleford KA, Moratorio G, et al. Low-Fidelity Polymerases of Alphaviruses Recombine at Higher Rates To Overproduce Defective Interfering Particles. J Virol. 2015;90(5):2446–54. doi:10.1128/JVI.02921-15. PubMed PMID:26676773; PubMed Central PMCID:PMCPMC4810721.

18. Xiao CT, Liu ZH, Yu XL, Ge M, Li RC, Xiao BR, et al. Identification of new defective interfering RNA species associated with porcine reproductive and respiratory syndrome virus infection. Virus Res. 2011;158(1-2):33–6. Epub 2011/03/10. doi:10.1016/j.virusres.2011.03.001. PubMed PMID:21385595.

19. Odagiri T, Tashiro M. Segment-specific noncoding sequences of the influenza virus genome RNA are involved in the specific competition between defective interfering RNA and its progenitor RNA segment at the virion assembly step. J Virol. 1997;71(3):2138–45. PubMed PMID:9032347; PubMed Central PMCID:PMCPMC191316.

20. Shingai M, Ebihara T, Begum NA, Kato A, Honma T, Matsumoto K, et al. Differential type I IFN-inducing abilities of wild-type versus vaccine strains of measles virus. J Immunol. 2007;179(9):6123–33. Epub 2007/10/20. doi:179/9/6123 [pii]. PubMed PMID:17947687.

21. Ho TH, Kew C, Lui PY, Chan CP, Satoh T, Akira S, et al. PACT- and RIG-I-Dependent Activation of Type I Interferon Production by a Defective Interfering RNA Derived from Measles Virus Vaccine. J Virol. 2016;90(3):1557–68. doi:10.1128/JVI.02161-15. PubMed PMID:26608320; PubMed Central PMCID:PMCPMC4719617.

22. Leppert M, Kort L, Kolakofsky D. Further characterization of Sendai virus DI-RNAs: a model for their generation. Cell. 1977;12(2):539–52. Epub 1977/10/01. doi:0092-8674(77)90130-1 [pii]. PubMed PMID:199355.

23. Perrault J. Origin and replicaition of defective interfering particles. In: Shatkin AJ, editor. Initiation Signal in Viral Gene Expression. Berlin: Springer-Verlag; 1981. p. 151–207.

24. Fearns R, Collins PL. Role of the M2-1 transcription antitermination protein of respiratory syncytial virus in sequential transcription. J Virol. 1999;73(7):5852–64. PubMed PMID:10364337; PubMed Central PMCID:PMCPMC112646.

25. Shcherbo D, Murphy CS, Ermakova GV, Solovieva EA, Chepurnykh TV, Shcheglov AS, et al. Farred fluorescent tags for protein imaging in living tissues. Biochem J. 2009;418(3):567–74. doi:10.1042/BJ20081949. PubMed PMID:19143658; PubMed Central PMCID:PMCPMC2893397.

26. Pfaller CK, Mastorakos GM, Matchett WE, Ma X, Samuel CE, Cattaneo R. Measles Virus Defective Interfering RNAs Are Generated Frequently and Early in the Absence of C Protein and Can Be Destabilized by Adenosine Deaminase Acting on RNA-1-Like Hypermutations. J Virol. 2015;89(15):7735–47. Epub 2015/05/15. doi:10.1128/jvi.01017-15. PubMed PMID:25972541; PubMed Central PMCID:PMCPmc4505647.

27. Yoshida A, Kawabata R, Honda T, Sakai K, Ami Y, Sakaguchi T, et al. A Single Amino Acid Substitution within the Paramyxovirus Sendai Virus Nucleoprotein Is a Critical Determinant for Production of Interferon-Beta-Inducing Copyback-Type Defective Interfering Genomes. J Virol. 2018;92(5). doi:10.1128/JVI.02094-17. PubMed PMID:29237838; PubMed Central PMCID:PMCPMC5809723.

28. Fearns R, Plemper RK. Polymerases of paramyxoviruses and pneumoviruses. Virus Res. 2017;234:87–102. doi:10.1016/j.virusres.2017.01.008. PubMed PMID:28104450; PubMed Central PMCID:PMCPMC5476513.

29. Kuo L, Fearns R, Collins PL. Analysis of the gene start and gene end signals of human respiratory syncytial virus: quasi-templated initiation at position 1 of the encoded mRNA. J Virol. 1997;71(7):4944–53. PubMed PMID:9188557; PubMed Central PMCID:PMCPMC191725.

30. Calain P, Roux L. The rule of six, a basic feature for efficient replication of Sendai virus defective interfering RNA. J Virol. 1993;67(8):4822–30. Epub 1993/08/01. PubMed PMID:8392616; PubMed Central PMCID:PMC237869.

31. Kolakofsky D, Pelet T, Garcin D, Hausmann S, Curran J, Roux L. Paramyxovirus RNA synthesis and the requirement for hexamer genome length: the rule of six revisited. J Virol. 1998;72(2):891–9. PubMed PMID:9444980.

32. Tawar RG, Duquerroy S, Vonrhein C, Varela PF, Damier-Piolle L, Castagne N, et al. Crystal structure of a nucleocapsid-like nucleoprotein-RNA complex of respiratory syncytial virus. Science (New York, NY). 2009;326(5957):1279–83. doi:10.1126/science.1177634. PubMed PMID:19965480.

33. Samal SK, Collins PL. RNA replication by a respiratory syncytial virus RNA analog does not obey the rule of six and retains a nonviral trinucleotide extension at the leader end. J Virol. 1996;70(8):5075–82. PubMed PMID:8764015; PubMed Central PMCID:PMCPMC190462.

34. Fodor E, Mingay LJ, Crow M, Deng T, Brownlee GG. A single amino acid mutation in the PA subunit of the influenza virus RNA polymerase promotes the generation of defective interfering RNAs. J Virol. 2003;77(8):5017–20. PubMed PMID:12663810; PubMed Central PMCID:PMCPMC152145.

35. Odagiri T, Tobita K. Mutation in NS2, a nonstructural protein of influenza A virus, extragenically causes aberrant replication and expression of the PA gene and leads to generation of defective interfering particles. Proc Natl Acad Sci U S A. 1990;87(15):5988–92. PubMed PMID:2143025; PubMed Central PMCID:PMCPMC54455.

36. Kang CY, Weide LG, Tischfield JA. Suppression of vesicular stomatitis virus defective intefering particle generation by a function(s) associated with human chromosome 16. J Virol. 1981;40(3):946–52. PubMed PMID:6275129; PubMed Central PMCID:PMCPMC256708.

37. Whistler T, Bellini WJ, Rota PA. Generation of defective interfering particles by two vaccine strains of measles virus. Virology. 1996;220(2):480–4. PubMed PMID:8661398.

38. Kingsbury DW, Portner A. On the genesis of incomplete Sendai virions. Virology. 1970;42(4):872–9. Epub 1970/12/01. PubMed PMID:4321309.

39. Yount JS, Kraus TA, Horvath CM, Moran TM, Lopez CB. A novel role for viral-defective interfering particles in enhancing dendritic cell maturation. J Immunol. 2006;177(7):4503–13. PubMed PMID:16982887.

40. Sun Y, López CB. Preparation of Respiratory Syncytial Virus with High or Low Content of Defective Viral Particles and Their Purification from Viral Stocks. Bio-protocol. 2016;6(10). Epub 5/20/2016. doi:0.21769/BioProtoc.1820.

41. Hotard AL, Shaikh FY, Lee S, Yan D, Teng MN, Plemper RK, et al. A stabilized respiratory syncytial virus reverse genetics system amenable to recombination-mediated mutagenesis. Virology. 2012;434(1):129–36. doi:10.1016/j.virol.2012.09.022. PubMed PMID:23062737; PubMed Central PMCID:PMCPMC3492879.

42. Zhao L, Jha BK, Wu A, Elliott R, Ziebuhr J, Gorbalenya AE, et al. Antagonism of the interferon-induced OAS-RNase L pathway by murine coronavirus ns2 protein is required for virus replication and liver pathology. Cell Host Microbe. 2012;11(6):607–16. doi:10.1016/j.chom.2012.04.011. PubMed PMID:22704621; PubMed Central PMCID:PMCPMC3377938.

43. Xu J, Sun Y, Li Y, Ruthel G, Weiss SR, Raj A, et al. Replication defective viral genomes exploit a cellular pro-survival mechanism to establish paramyxovirus persistence. Nat Commun. 2017;8(1):799. doi:10.1038/s41467-017-00909-6. PubMed PMID:28986577; PubMed Central PMCID:PMCPMC5630589.

44. Wickham H. ggplot2: Elegant Graphics for Data Analysis. New York: Springer-Verlag; 2016.

45. Langmead B, Salzberg SL. Fast gapped-read alignment with Bowtie 2. Nat Methods. 2012;9(4):357–9. doi:10.1038/nmeth.1923. PubMed PMID:22388286; PubMed Central PMCID:PMCPMC3322381.

