## Supplemental Information for "A specific sequence in the genome of respiratory syncytial virus regulates the generation of copy-back defective viral genomes"

**Supplementary Information**

**SI Figures:**

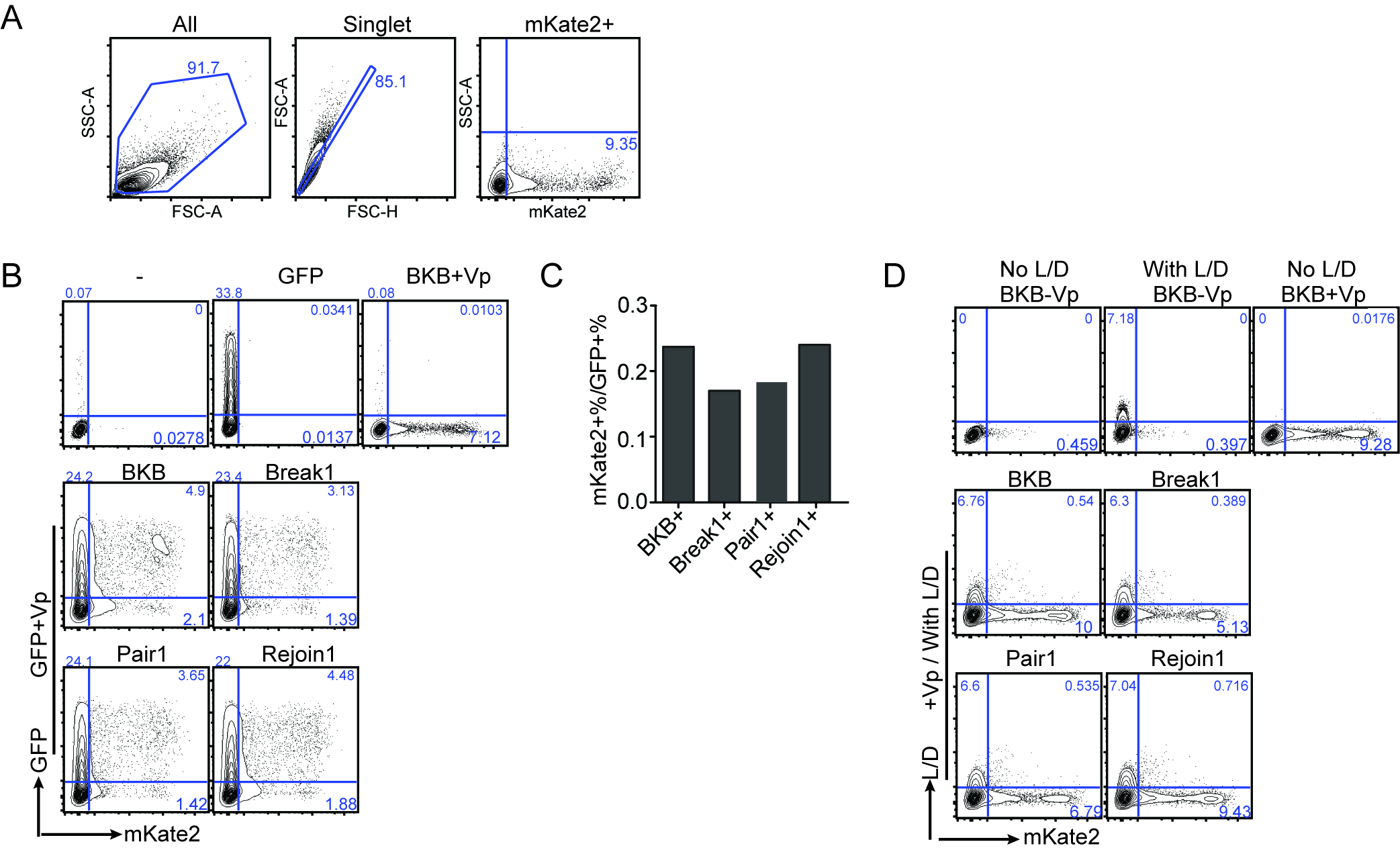

**S1 Fig.** Identification of RSV genomic regions responsible for the generation of cbDVGs. (A) Gating strategy for all flow cytometry data shown in Fig. 2, 4A and S3 (B-C) BSR-T7 cells were co-transfected with 4 helper plasmids expressing the polymerase proteins, pmax-GFP, and the designated minigenome construct. mKate2 expression and GFP expression were measured by flow cytometry. Representative flow plots are shown in (B) and fold change in (C). (D) BSR-T7 cells were co-transfected with the four helper plasmids and the designated minigenome construct. Cells were first stained with Live/Dead aqua followed by flow cytometry. Representative flow plots are shown.

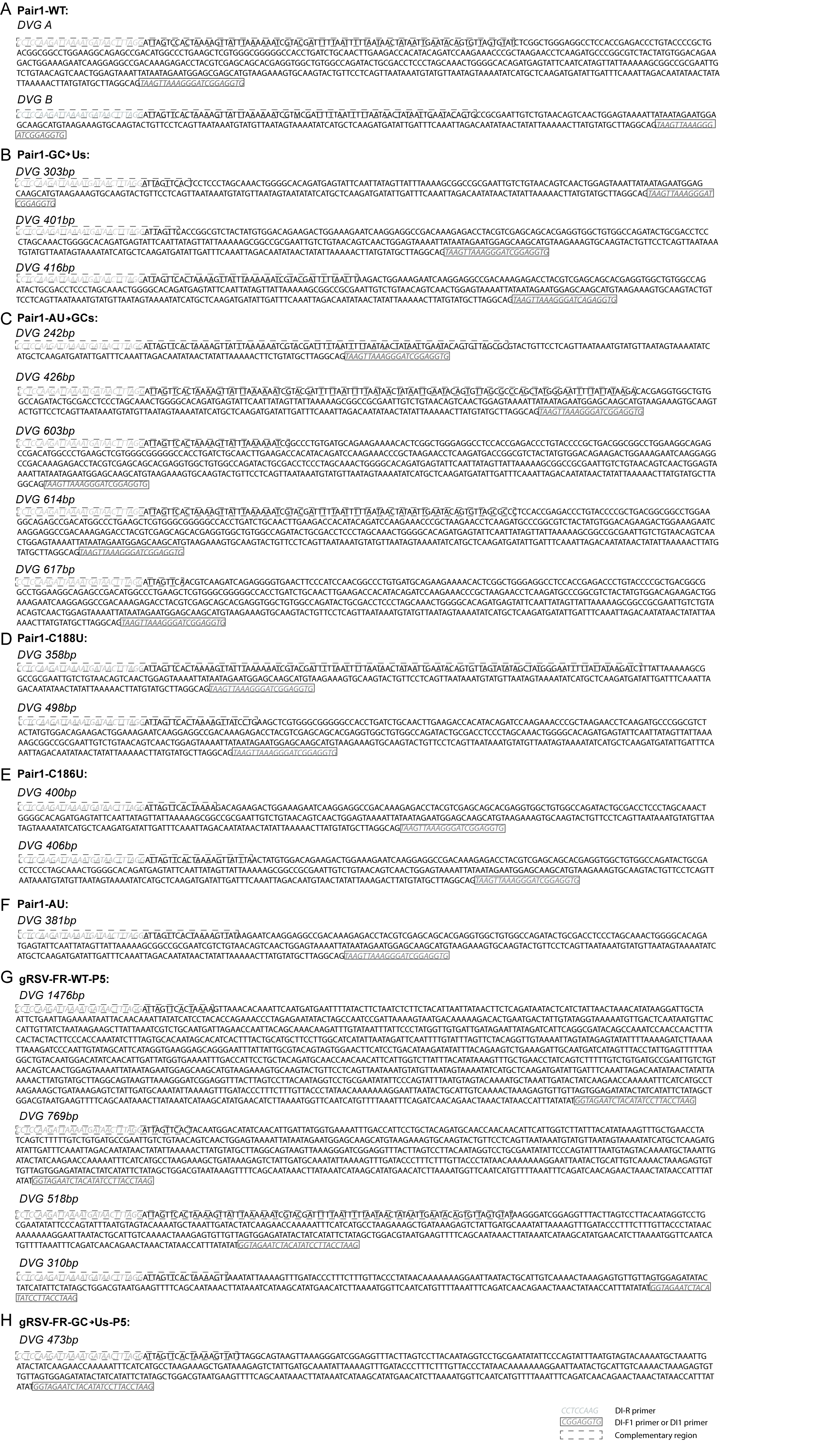

**S2 Fig.** Sequences of cbDVGs identified from labeled bands in RSV minigenome system (A-F), and in gRSV-FR-WT (G) or gRSV-FR-GC >Us viral infections (H).

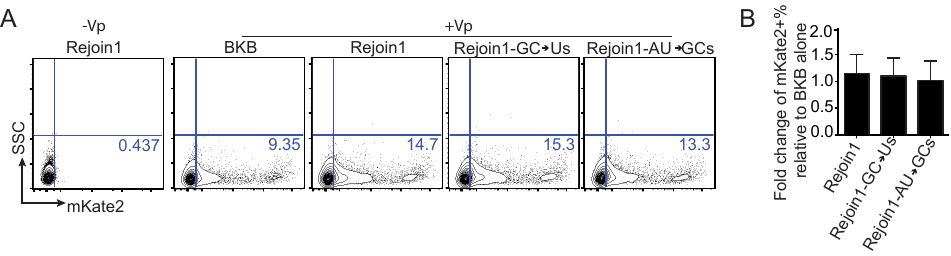

**S3 Fig.** GC mutations in the RSV minigenome did not alter viral polymerase processivity. The same mutations contained in the Pair1 construct were introduced in the Rejoin1 construct. BSR-T7 cells were co-transfected with all 4 helper plasmids as well as BKB or Rejoin1 mutations. mKate2 expression was measured by flow cytometry. Representative flow plots are shown in panel (A), and quantification of three repeats is shown as fold change in (B). Fold change was calculated as the percentage of mKate2 expressing cells transfected with Rejoin1 or its two mutants over BKB controls.

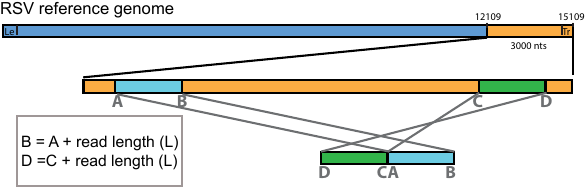

**S4 Fig.** Schematic representation of how VODKA constructs a theoretic library for cbDVGs from the last 3000 nucleotides (nts) of the genome.

**S1 Table. Oligo primer list.**

| **Primer name** | **Sequences (5’-3’)** | **Application in paper** |
| --- | --- | --- |
| DI1 (RSV) | CTTAGGTAAGGATATGTAGATTCTACC | DVG-RT-PCR |
| DI-R | CCTCCAAGATTAAAATGATAACTTTAGG | DVG-RT-PCR |
| SeV DI1 | GGTGAGGAATCTATACGTTATAC | DVG-RT-PCR |
| gSeV DI1 | ACCAGACAAGAGTTTAAGAGATATGTATT | DVG-RT-PCR |
| DI-F1 | CACCTCCGATCCCTTTAACTTA | DVG-RT-PCR |
| DI-F2 | CAATATCATCTTGAGCATGATATTTTAC | DVG-RT-PCR |
| DI-F3 | CATTATTCATTATGAAAGTTGTATAACAGACTAC | DVG-RT-PCR |
| RSV G-F | AACATACCTGACCCAGAATC | qPCR |
| RSV G-R | GGTCTTGACTGTTGTAGATTGCA | qPCR |
| *GAPDH-F* | CTCCCACTCTTCCACCTTCG | qPCR |
| *GAPDH-R* | CCACCACCCTGTTGCTGTAG | qPCR |
| T7 term-R | CCGG*GAATTC*AACATATAGTTCCTCCTTTCAGCA | Cloning |
| RSV TrC37-F | GCGC*ACTAGT*GACCT*GGGTCCC*TTAG*GCTCTTC*TTAAAAATCGTACGATTTTTTAAATAACTTTTAGTGAAC | Cloning |
| T7 prom-F | GCGC*ACTAGT*TAATACGACTCACTATAGGGAAGTTT | Cloning |
| mKateGE-R | CCG*GGGACCC*CTAA*GCGGCCGC*TTTTTAATAACTATAATTGAATACTCATCTGTGCC | Cloning |
| TrC+Rejoin1-F | GCGCA*GGGTCCC*TTAGG*CCATGG*AGCTAG*GCTCTTC*AAAACTGATTAAAATCACAGGTAGTCTGTTATAC | Cloning |
| Break1-F | GGCCA*GCGGCCGC*GAATTGTCTGTAACAGTCAACTGG | Cloning |
| Break1-R | CGCGA*GGGTCCC*ACCTCCGATCCCTTTAACTTACT | Cloning |
| Rejoin1-F | GGCCA*GGGTCCC*AAAACTGATTAAAATCACAGGTAGTCTG | Cloning |
| Rejoin1-R | CGCGA*GCTCTTC*TTTAATTTTTAATAACTATAATTGAATACAGTGTTAGTG | Cloning |
| GCs->Us-F | AAAAATCGTACGATTTTTAATTTTTAATAACTATAATTGAATACAGTGTTAAAAAAAAGCTATGGGAATTTTTATTATAAGATCTTTATTCATTATTCATTATGAAAG | Site-mutagenesis |
| GCs->Us-R | CTTTCATAATGAATAATGAATAAAGATCTTATAATAAAAATTCCCATAGCTTTTTTTTAACACTGTATTCAATTATAGTTATTAAAAATTAAAAATCGTACGATTTTT | Site-mutagenesis |
| AU->GCs-F | GTACGATTTTTAATTTTTAATAACTATAATTGAATACAGTGTTAGCGCCCAGCTATGGGAATTTTTATTATAAGATCTTTATTCATTATTCATTA | Site-mutagenesis |
| AU->GCs-R | TAATGAATAATGAATAAAGATCTTATAATAAAAATTCCCATAGCTGGGCGCTAACACTGTATTCAATTATAGTTATTAAAAATTAAAAATCGTAC | Site-mutagenesis |
| C1284T-F | GAATAAAGATCTTATAATAAAAATTCCCATAGCTATATACTAACACTGTATTCAATTATAGTTATTAAAAA | Site-mutagenesis |
| C1284T-R | TTTTTAATAACTATAATTGAATACAGTGTTAGTATATAGCTATGGGAATTTTTATTATAAGATCTTTATTC | Site-mutagenesis |
| C1286T-F | TTAATTTTTAATAACTATAATTGAATACAGTGTTAATGTATAGCTATGGGAATTTTTATTATAAGATCTTT | Site-mutagenesis |
| C1286T-R | AAAGATCTTATAATAAAAATTCCCATAGCTATACATTAACACTGTATTCAATTATAGTTATTAAAAATTAA | Site-mutagenesis |
| mut virus PmlI fwd | TTAACACGTGGTGAGAGAGGACCCACTAAA | Site-mutagenesis for virus |
| mut virus MulI rev | GCTTACGCGTATATAGTTCCTCCTTTCAGCAAAA | Site-mutagenesis for virus |
| A mut virus rev | CTATAATTGAATACAGTGTTAAAAAAAAGCTATGGGAATTTTTATTATAAGATC | Site-mutagenesis for virus |
| A mut virus fwd | GATCTTATAATAAAAATTCCCATAGCTTTTTTTTAACACTGTATTCAATTATAG | Site-mutagenesis for virus |
